# Single-cell transcriptome and T cell receptor profiling of the tuberculin skin test

**DOI:** 10.1101/2024.06.25.600676

**Authors:** Carolin T. Turner, Joshua Rosenheim, Clare Thakker, Aneesh Chandran, Holly Wilson, Cristina Venturini, Gabriele Pollara, Benjamin M. Chain, Gillian S. Tomlinson, Mahdad Noursadeghi

## Abstract

The tuberculin skin test (TST) is a cutaneous delayed hypersensitivity reaction to antigen from *Mycobacterium tuberculosis* (Mtb). We provide the first single cell sequencing characterisation of the human TST reaction, based on skin suction blisters induced at the site of the TST on day 2 in 31 individuals. Integrated single cell RNA and TCR sequencing showed the immune response to be dominated by T cells, with smaller populations of NK cells and myeloid cells. T cells comprised CD4, CD8, gamma/delta and NK T cells, with 50% of all T cells identified as cytotoxic and 14% as regulatory. Interferon gamma gene expression was strongest in CD8 T cells, and distinct CD4 T helper lineages could not unambiguously be identified at this time point. Amongst myeloid cells, 63% displayed antimicrobial gene expression and 28% were functionally polarised towards antigen presentation with higher levels of HLA class 2 expression. We derived and validated transcriptional signatures for cell types and cellular functions relevant to the immune landscape of the TST. These data help to improve our understanding of the immune response to Mtb and enable further exploration of bulk transcriptomic data through context-specific cellular deconvolution.

## Introduction

The tuberculin skin test (TST) is a cutaneous delayed hypersensitivity reaction to the recall antigen purified protein derivative (PPD) from *Mycobacterium tuberculosis* (Mtb) and is used clinically as a measure of T cell memory for Mtb antigens. Employing the TST as a standardised human in vivo challenge model, we have previously shown that genome-wide transcriptional profiling of skin punch biopsies from the site of the TST allows highly sensitive and comprehensive assessments of the complex immune response to Mtb antigens that recapitulates the transcriptional perturbations occurring at the site of human tuberculosis (TB) disease (1–4).

The current understanding of the TST reaction (5–7) involves infiltration of innate immune cells in response to skin injury and the presence of antigen. Antigen is taken up and processed by myeloid cells that subsequently migrate to the lymph node for antigen presentation to circulating Mtb-reactive memory T cells. T cells then migrate to the TST site and amplify the inflammatory response through cytokine production, particularly interferon gamma (IFNG) (1,8–10), resulting in skin erythema and induration, which is maximal at days 2-3. The necessity for memory T cells in this amplification process is demonstrated by the fact that individuals with no previous exposure to mycobacterial antigen do not mount a clinically measurable TST response (1). Furthermore, it is known that CD4 T cells are required for a functional TST response, as individuals with HIV and low CD4 T cell counts frequently show reduced or negative TST results (3,11).

To evaluate the cell composition of multicellular tissue transcriptomes, multiparameter gene signatures have been proposed to reflect distinct cell types or functional responses (12,13). Applying such gene signatures to bulk transcriptional data from TST biopsies taken on day 2 shows selective accumulation of T cells, monocyte-derived and natural killer cells (2–4), in accordance with early histological data (14). Skin suction blisters are a less invasive alternative to skin punch biopsies, whereby application of negative pressure to the surface of the skin results in cell migration into the blister fluid, thus allowing sampling of single cells from tissue without the need for complex dissociation steps (15). Flow cytometric analyses of skin suction blisters from the site of the TST have yielded valuable insights into the kinetics of the T cell dominated immune response, focussing on memory and regulatory CD4 T cells (16–18). A full characterisation of the cellular complexity by single cell analysis of the TST has so far only been attempted in the guinea pig model (19).

Here, we apply single cell RNA and TCR sequencing to human TST suction blisters to characterise the cellular composition of the day 2 TST and enable annotation of cell types with functional attributes. In addition, we derive transcriptional signatures for cell types and cellular functions relevant to the immune landscape of the TST. These data help to improve our understanding of the immune response in TB and enable further exploration of bulk transcriptomic data through context-specific cellular deconvolution.

## Results

### Cell types present in day 2 TST suction blisters

We undertook single cell RNA, TCR and antibody-derived tag (ADT) sequencing of day 2 TST suction blister cells from 31 individuals (**Table 1**) with immunological memory to Mtb antigens. Following quality control filtering and data integration, this resulted in expression data for 18291 genes and 130 surface proteins in 63881 cells, with concurrent TCR sequencing data for 37413 cells (**Table S1**). To identify cell types and subsets, we performed sequential clustering of the cells, based on similarity of gene expression profiles. We applied Louvain clustering and confirmed statistical validity of the identified clusters in each sub-clustering round by post-hoc significance analysis (20). The number of digits in the cluster label reflects the number of sub-clustering steps performed to obtain the final cluster, with the digits referring to the cluster assignment in each round (**Table S2**). This approach enabled finer separation of distinct cell populations that were initially grouped together, such as melanocytes and keratinocytes in cluster 1, CD4 and CD8 T cells present in clusters 2 and 3, natural killer (NK) and NKT cells in cluster 4, or gamma/delta (gd) and CD8 T cells in clusters 5 and 8 (**Figure S1**). In total, we resolved 101 clusters, for which we sought ontological labels using independently established canonical gene or protein markers of cell types relevant to the TST (**Figure 1A**, **Table S3**, **Table S4**).

**Figure 1.**
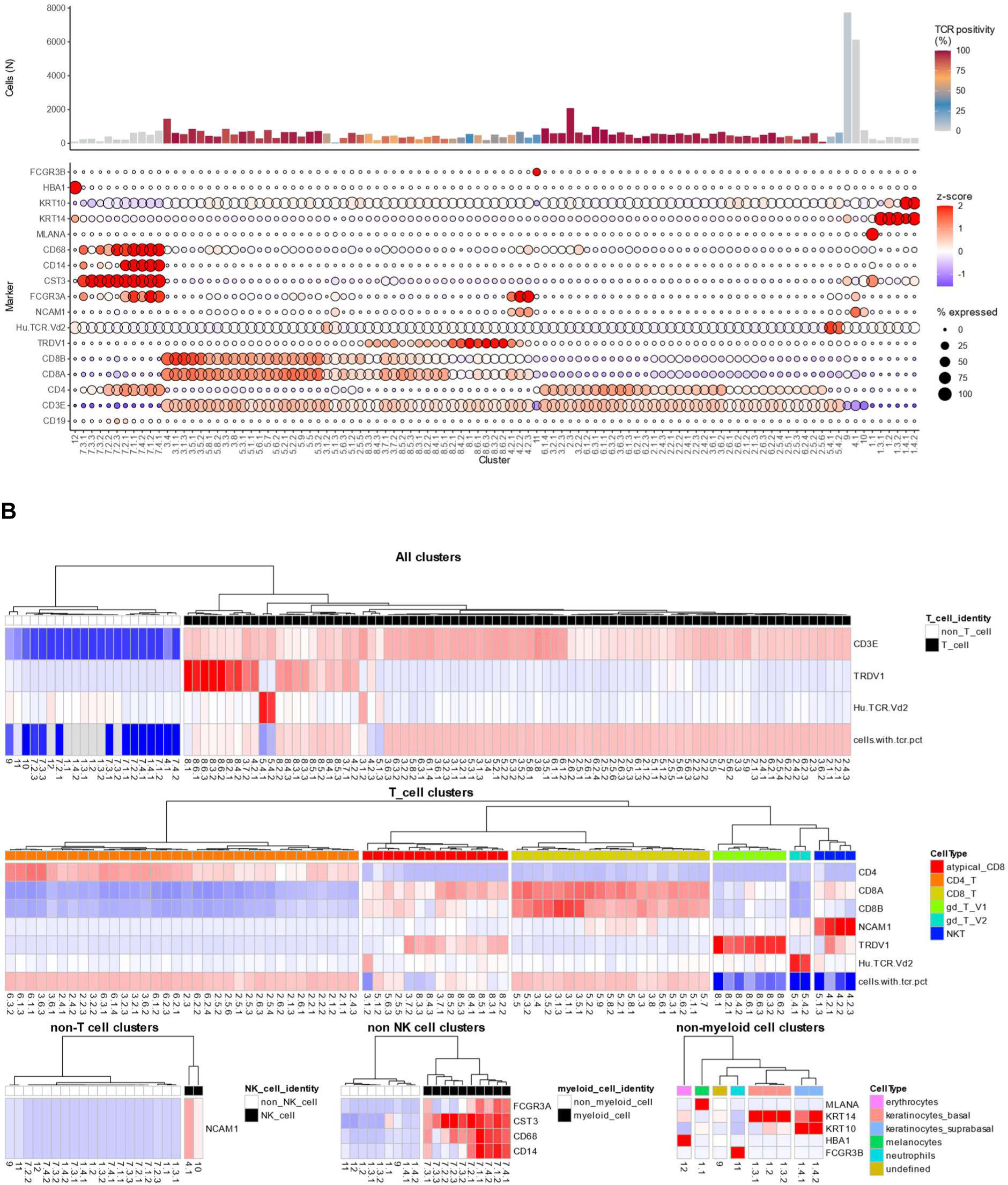
Cell types present in suction blisters at the site of the tuberculin skin test on day 2. **A.** Number of cells, percentage of cells with single cell TCR sequencing data, and expression of canonical gene or protein markers for known cell types in each cluster. In the bar graph, bar height represents the cluster size as shown on the y-axis, and colour indicates the percentage of TCR positivity for each cluster. In the dot plot, dot size represents the percentage of cells expressing each marker in each cluster, and colour shows the Z-score scaled expression of the marker, calculated compared to all other cells in the dataset, and averaged for each cluster. The Z-score colour scale is capped at -2 and 2. Protein markers are prefixed with ‘Hu’. Clusters were ordered along the x-axis by hierarchical clustering (ward D2 method) with the set of markers shown on the y-axis. **B.** Heatmap representations of the Z-score scaled expression of different subsets of markers from A in different subsets of clusters. Z-scores were calculated across all cells in the dataset to define T cell vs. non-T cell clusters (top row), across all T cells to define different T cell subsets (middle row), or across all non-T cells to define any other cell types (bottom row). The Z-score colour scale is the same as in A. The dendrograms for each heatmap show hierarchical clustering (ward D2 method) of the selected cell clusters based on the chosen canonical gene and protein markers, with the colour legends indicating the resulting annotation of the clusters as different cell types.

**Table 1.** Demographic data summary. * Participants of ‘other’ ethnicity were born in the Philippines (n=4), Romania (n=1), Syria (n=1) or Iraq (n=1). ** Induration measurements were recorded for 30 out of 31 participants.

|  |  | Number | Percentage |
| --- | --- | --- | --- |
| **Total** |  | 31 | 100 |
| **Sex** | Male | 9 | 29 |
|  | Female | 22 | 71 |
| **Median age in years (range)** |  | 33 (19-48) |  |
| **Ethnicity** | Black African | 16 | 51.6 |
|  | Indian | 4 | 12.9 |
|  | White | 4 | 12.9 |
|  | Other * | 7 | 22.6 |
| **Median injection site induration in mm (range) \*\*** |  | 14 (7-26) |  |

Approximately two thirds of all cells were identified as T cells (expressing CD3E and either a high percentage of alpha/beta TCR sequencing data or markers for Vδ1 and Vδ2 gd T cells) (**Figure 1A-B**). These were further separated into CD4 T cells (30% of all cells, 45% of T cells), two populations of CD8 T cells (together making up 28% of all cells and 42% of T cells), as well as smaller populations of Vδ1 gd T cells (4.3% of all cells, 6.4% of T cells), Vδ2 gd T cells (1.6% of all cells, 2.4% of T cells) and NKT cells (2.5% of all cells, 3.7% of T cells) (**Figure 1A-B**, **Table 2**, **Table S4**). We annotated the two CD8 T cell populations as ‘classical’ (19.4% of all cells, 29% of T cells) and ‘atypical’ CD8 T cells (8.9% of all cells, 13.4% of T cells). ‘Atypical’ CD8 T cells exhibited lower expression of CD8A and CD8B, and a smaller percentage in which we were able to detect alpha/beta TCR sequences.

**Table 2.** Summary of cell types and functions identified in suction blisters at the site of the tuberculin skin test on day 2. The ‘parent’ cell type is either T_cell for T cell subsets and T cell function, or Myeloid for myeloid cell function. TCR = T cell receptor. N/A = no signature derivation attempted.

|  | Number clusters | Number cells | % of all cells | % of 'parent' cell type | % TCR+ | Number signature genes |
| --- | --- | --- | --- | --- | --- | --- |
| Full dataset | 101 | 63881 | 100 |  | 58.57 | N/A |
| **ONTOGENY** |  |  |  |  |  |  |
| T_cell | 80 | 42746 | 66.92 |  | 86.27 | 50 |
| NK | 2 | 6911 | 10.82 |  | 0.33 | 50 |
| Myeloid | 10 | 4046 | 6.33 |  | 0.69 | 50 |
| Neutrophil | 1 | 513 | 0.80 |  | 27.88 | N/A |
| Keratinocyte_basal | 3 | 919 | 1.44 |  | 0.00 | N/A |
| Keratinocyte_suprabasal | 2 | 626 | 0.98 |  | 0.16 | N/A |
| Melanocyte | 1 | 266 | 0.42 |  | 0.00 | N/A |
| Erythrocyte | 1 | 116 | 0.18 |  | 0.00 | N/A |
| Undefined | 1 | 7738 | 12.11 |  | 4.39 | N/A |
| **T CELL SUBSETS** |  |  |  |  |  |  |
| CD4_T | 34 | 19287 | 30.19 | 45.12 | 97.31 | 13 |
| CD8_T | 19 | 12394 | 19.40 | 28.99 | 94.64 | 9 |
| atypical_CD8 | 14 | 5709 | 8.94 | 13.36 | 75.74 | 0 |
| gd_T_V1 | 7 | 2739 | 4.29 | 6.41 | 46.04 | 9 |
| gd_T_V2 | 2 | 1041 | 1.63 | 2.44 | 13.35 | 21 |
| NKT | 4 | 1576 | 2.47 | 3.69 | 41.62 | 34 |
| **T CELL FUNCTION** |  |  |  |  |  |  |
| T1 | 27 | 13368 | 20.93 | 31.27 | 77.86 | 9 |
| T2 | 13 | 6399 | 10.02 | 14.97 | 96.58 | 0 |
| T22 | 9 | 6002 | 9.40 | 14.04 | 97.05 | 1 |
| Treg | 10 | 5982 | 9.36 | 13.99 | 98.51 | 50 |
| Naive | 16 | 6047 | 9.47 | 14.15 | 62.89 | 9 |
| Cytotoxic | 34 | 21291 | 33.33 | 49.81 | 85.04 | 10 |
| **MYELOID SUBSETS** |  |  |  |  |  |  |
| Macrophage | 4 | 2557 | 4.00 | 63.20 | 0.31 | N/A |
| Classical DC | 2 | 632 | 0.99 | 15.62 | 0.79 | N/A |
| mregDC | 1 | 267 | 0.42 | 6.60 | 2.25 | N/A |
| pDC | 1 | 222 | 0.35 | 5.49 | 2.70 | N/A |
| Langerhans cell | 1 | 123 | 0.19 | 3.04 | 0.00 | N/A |
| **MYELOID CELL FUNCTION** |  |  |  |  |  |  |
| Ag_presenting | 4 | 1121 | 1.75 | 27.71 | 1.52 | 50 |
| Antimicrobial | 4 | 2557 | 4.00 | 63.20 | 0.31 | 50 |

Non-T cell clusters were identified as NK cells (expressing NCAM1; 10.8% of all cells), myeloid cells (CST3+; 6.3%), neutrophils (FCGRB+; 0.8%), two populations of keratinocytes (together 2.4%), melanocytes (MLANA+; 0.4%) and erythrocytes (HBA1+; 0.2%). One cluster remained undefined (cluster 9; 12.1%) (**Figure 1A-B**, **Table 2**, **Table S4**). The two keratinocyte populations represented basal keratinocytes, expressing KRT14, and suprabasal keratinocytes, additionally expressing the early differentiation keratin KRT10 (21,22). The cluster that could not be assigned a distinct cell type based on the selected gene and protein markers also did not yield any cluster-specific marker genes when interrogated for differential gene expression compared to other cell types by Wilcoxon rank sum test (**Table S5**). Instead, cells belonging to this cluster were characterised by a relatively smaller number of detected genes and total counts (**Figure S2**), and likely represent low-quality cells despite passing quality control filtering. We did not find any evidence of B cells in day 2 TST blisters.

### Functional T cell sub-types present in day 2 TST suction blisters

Next, we sought to discriminate the T cell cluster by gene expression that reflected functional attributes. We hypothesised that the day 2 TST T cell response is limited to activated memory T cells, comprising CD4 T cells of the Th1, Th17 and Treg lineages as well as cytotoxic CD8 T cells, but only low frequency of proliferating cells due to the early time point. We therefore included gene or protein markers for proliferation, cytotoxicity, T cell lineage, multifunctional cytokines, regulatory T cells, activation, and differentiation (**Figure 2A**, **Table S3**). Almost all T cell clusters showed evidence of activation, assessed by gene expression of CD40LG, CD69 or LAG3, whilst only a few clusters had enriched expression of proliferation markers (MKI67, PCLAF, CDK1) (**Figure 2A**). The proliferating cells were mainly evident among non-CD4 T cell clusters at this time point. These included clusters 5.1.2 (atypical CD8), 5.1.3 (NKT), 5.1.1 and 3.3 (both ‘classical’ CD8 T cells). A large proportion of T cell clusters expressed cytotoxic genes (GZMA, GZMB, GNLY, PRF1). In fact, all clusters annotated as ‘classical’ CD8 T cells, NKT cells or Vδ2 gd T cells were found to express marker genes for cytotoxic function. In addition, a subset of CD4 and ‘atypical’ CD8 T cell clusters were also associated with cytotoxic function (**Figure 2B**). Surprisingly, we found a clear group of T cell clusters with naive phenotype, identified by surface protein expression of CD45RA and CD27. These naive T cells mapped exclusively to Vδ1 gd T cells and atypical CD8 T cells (**Figure 2B**). Whilst there was an obvious set of CD4 Treg clusters (expressing FOXP3, CTLA4 and IL2RA), delineating other T cell lineages was undermined by low expression of lineage-specific cytokines and transcription factors (**Figure 2A**). IFNG was predominantly expressed by non-CD4 T cells, and IL17A and IL17F were absent from the dataset. We therefore defined T cell lineage irrespective of CD4/CD8 annotation as T1 (enriched for IFNG/TBX21), T2 (enriched for IL4/IL13/GATA3) or T22 (enriched for IL22/RORC) (**Figure 2B**).

**Figure 2.**
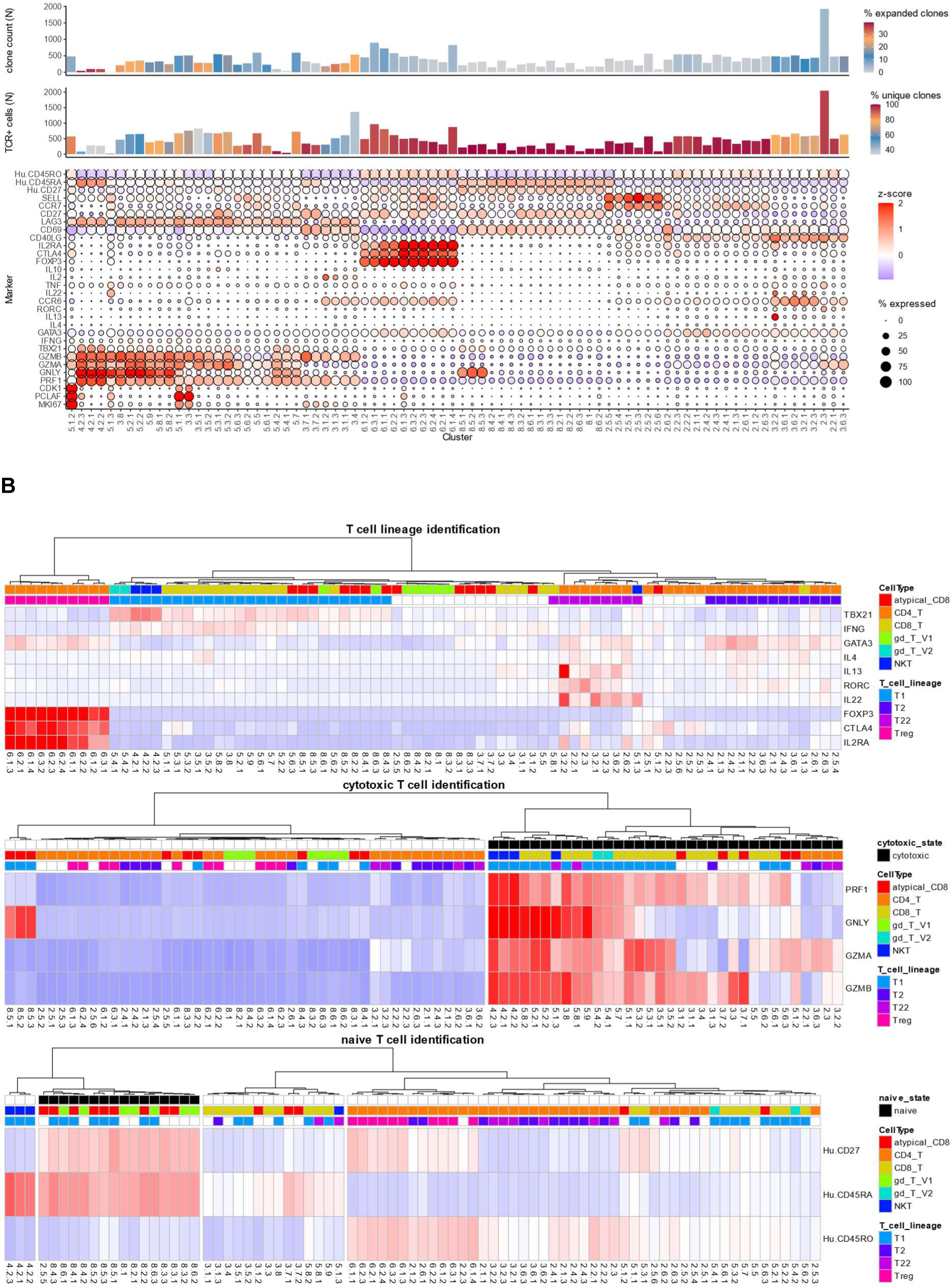
Functional T cell sub-types present in suction blisters at the site of the tuberculin skin test on day 2. **A.** Quantification of alpha/beta TCR pairs (clones) and expression of independently established gene or protein markers for known T cell function in each T cell cluster. For each cluster, the bar graphs represent on the y-axes the number of cells with single cell TCR sequencing data and the number of unique alpha/beta TCR clones, while the colour scales indicate the percentage of unique and expanded clones, respectively. Expanded clones are defined as TCR clones that are found more than once. In the dot plot, dot size represents the percentage of cells expressing each marker in each cluster, and colour shows the Z-score scaled expression of the marker, calculated compared to all other T cells in the dataset, and averaged for each cluster. The Z-score colour scale is capped at -2 and 2. Protein markers are prefixed with ‘Hu’. Clusters were ordered along the x-axis by hierarchical clustering (ward D2 method) with the set of markers shown on the y-axis. **B.** Heatmap representations of the Z-score scaled expression of three different subsets of markers from A, with the same Z-score colour scale as in A. The dendrograms show hierarchical clustering (ward D2 method) of the T cell clusters based on the selected markers, to identify different T cell lineages (top), cytotoxic T cells (middle) and naive T cells (bottom). The colour legends indicate annotation of T cell clusters according to these clustering analyses (T cell lineage, cytotoxic state, and naive state), or as defined in Figure 1B (CellType).

Quantification of the different functional T cell subsets showed that 50% of all T cells were cytotoxic, 14% naive, and 14% regulatory (**Table 2**, **Table S4**, **Figure 2B**). These functions were non-overlapping. In addition, 31% of T cells were assigned to a T1 lineage, 15% to a T2 lineage, and 14% to a T22 lineage. Notably, none of the T1 clusters was annotated as CD4 T cells. Instead, the strongest IFNG and TBX21 signals were detected in CD8 T cells, and Vδ2 gd T or NKT cells, respectively (**Figure 2B**). The majority of T1 cells were also annotated as cytotoxic, with the rest assigned as naive. About half of the T22 clusters were also annotated as cytotoxic, whereas most T2 cells were not annotated with either of these functions.

Naive T cell clusters did not contain any TCR clones present more than once (**Figure 2A**). Similarly, Treg cell clusters were almost exclusively composed of unique TCR clones. In contrast, the highest proportions of expanded (>1) T cell clones were found in NKT clusters (4.2.3, 4.2.1, 4.2.2), followed by other cytotoxic T cell clusters. Of note, enriched expression of proliferation markers in T cell clusters did not correlate with proportion of expanded TCR clones at this time point.

### Myeloid cell subsets present in day 2 TST suction blisters

To assess the ontological and functional heterogeneity of myeloid cells in the TST, we used gene markers for monocyte, macrophage, and dendritic cell (DC) populations, as well as for cytokines associated with canonical pro-inflammatory or regulatory roles, or induction of distinct T cell subsets, and genes involved with antimicrobial, and antigen processing or presentation pathways (**Figure 3A**, **Table S3**).

**Figure 3.**
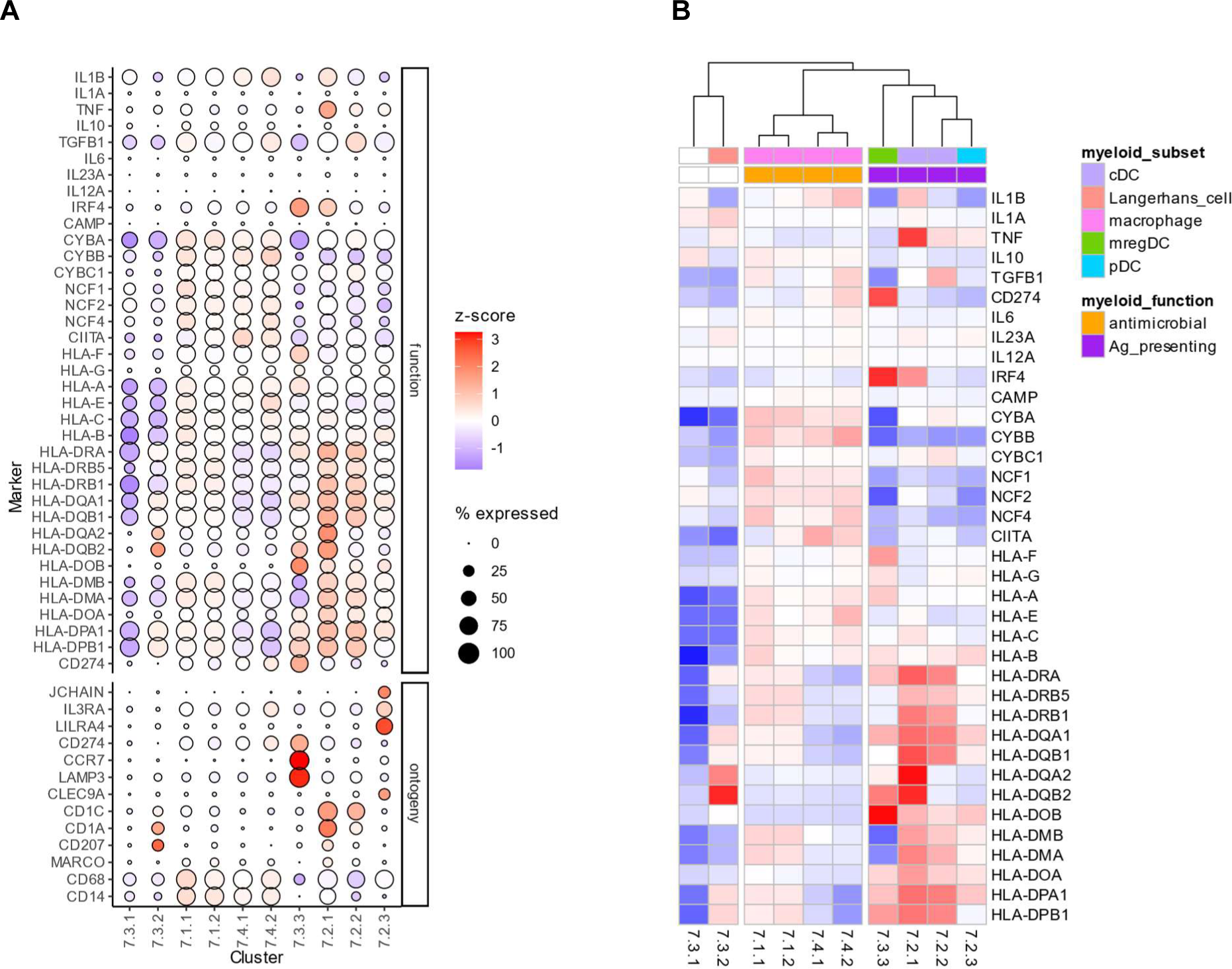
Myeloid cell subsets present in suction blisters at the site of the tuberculin skin test on day 2. **A.** Dot plot visualisation of the expression of independently established marker genes for different functional and ontological myeloid subsets. Dot size represents the percentage of cells expressing the gene in each cluster, and colour shows the Z-score scaled gene expression, calculated compared to all other myeloid cells, and averaged for each cluster. The colour scale is capped at -2 and 2. Clusters were ordered along the x-axis by hierarchical clustering (ward D2 method) with the set of functional markers shown on the y-axis. **B.** Heatmap representation of a subset of marker genes from A. The dendrogram shows hierarchical clustering (ward D2 method) of the myeloid cells into three groups based on the Z-score scaled expression of marker genes for different myeloid functions (Z-score colour scale as in A.). No functional annotation assigned to myeloid clusters 7.3.1 and 7.3.2. Ag_presenting = antigen-presenting.

We were able to identify macrophages (expressing CD14 and CD68; 63.2% of myeloid cells), ‘classical’ DC (CD1C+; 15.6%), mature DCs enriched in immunoregulatory molecules (mregDC) (LAMP3+CCR7+CD274+; 6.6%), plasmacytoid DCs (pDCs) (LILRA4+IL3RA+JCHAIN+; 5.5%), and Langerhans cells (CD207+; 3%), with a further myeloid sub-population remaining undefined (cluster 7.3.1; 6%) (**Figure 3A**, **Table 2**, **Table S4**).

Using the functional markers, myeloid cells separated into subsets with antimicrobial or antigen-presenting function (**Figure 3B**), with enriched expression of either genes encoding the phagocyte oxidase complex or HLA class 2 proteins, respectively. In accordance with their known roles in a cellular immune response, myeloid clusters classified as macrophages displayed enriched antimicrobial function, whereas myeloid clusters identified as classical, plasmacytoid and mreg DCs exhibited enriched antigen-presenting function. We were unable to assign Langerhans cells and the undefined myeloid cluster a function in this dichotomous classification. Overall, 63% of myeloid cells were defined as antimicrobial and 28% were defined as antigen-presenting (**Table 2**, **Table S4**). Antigen-presenting DCs were also enriched for TNF expression, whereas other cytokines showed no differential expression and were often expressed by a very small percentage of myeloid cells (**Figure 3A**). In addition to a canonical role for type 2 IFN responses to Mtb, interest in type 1 IFN responses has also emerged (23). Expression of type 1 IFN genes were not detected in our dataset. However, we also probed for evidence of IFN activity using expression of multi-gene signatures specifically induced by either type 1 or type 2 IFNs (3). This approach provided evidence for both type 1 and type 2 IFN activity which co-segregated with distribution of their receptor expression across cell types (**Figure S3**). The type 2 IFN inducible signature was mainly limited to myeloid cells, whereas the type 1 IFN inducible signature was more widely distributed, albeit not ubiquitous across all cell types.

### TST blister gene signatures

We aimed to derive gene signatures for the cell types and functions identified in the day 2 TST blister, with the motivation to obtain context-specific signatures that can be applied in future work to deconvolute TST biopsy bulk transcriptomes collected from different disease groups or over time. To this end, we created ‘pseudobulk’ data by summing the gene counts of single cells belonging to the same sample/cluster label combination, and then assigned them to a binary class (cell type of interest versus any other cell type) for each desired signature. Cell type-specific signature genes were identified through repeated Wilcoxon tests of equal-sized subsamples for each cell type class, and average statistics across all iterations were used to select a maximum of 50 signature genes, based on over-expression in the cell type of interest.

We derived signatures for T cells, NK cells and myeloid cells, as well as for T cell subsets (CD4_T, CD8_T, atypical_CD8, gd_T_V1, gd_T_V2, NKT, T1, T2, T22, Treg, cytotoxic, and naive), and functional myeloid subsets (antimicrobial and antigen-presenting). **Table S6** summarises the identified signature genes for those cell types and functions. No significant signature genes were found for T2 and atypical CD8, suggesting that the corresponding clusters could not efficiently be discerned as separate cell type population. In addition, the T22 signature contained only one gene (CD40LG), potentially reducing its power to distinguish between distinct T cell subsets. Other signatures ranged from 9 to 50 genes (**Table 2**). As expected, based on the overlapping annotation of cytotoxic, T1 and CD8 T cells (**Figure 2B**), their signatures also shared several genes (e.g. CCL5, NKG7). Similarly, the naive and gd_T_V1 signatures shared several genes (e.g. BACH2, CD7).

To internally validate the derived signatures, we quantified their expression in each cell of the TST blister dataset, and averaged signature Z-scores across individual cluster labels (**Figure S4**), or across identified cell types and functional subsets (**Figure 4**). In general, each signature achieved highest average expression in its target cell type (**Figure 4A-D**). An exception to this was the CD8_T signature which showed similar enrichment in clusters annotated as CD8 T cells, NKT or gd T cells (**Figure 4A**, **Figure S4A**). Furthermore, the T1 and T22 signatures did not discriminate well between their ‘true’ target clusters and other CD8 or CD4 clusters, respectively (**Figure 4B**, **Figure S4B**). To summarise how well the signatures distinguished the target cell type from other cells, we calculated the area under the receiver operating curve (AUROC) for each signature after assigning each cell to a binary class according to their cluster annotation (target cell type or any other cell type). Most signatures achieved AUROC values of >90%, whereas the one-gene signature T22 yielded a lower AUROC value of <80% (**Table 3**).

**Figure 4.**
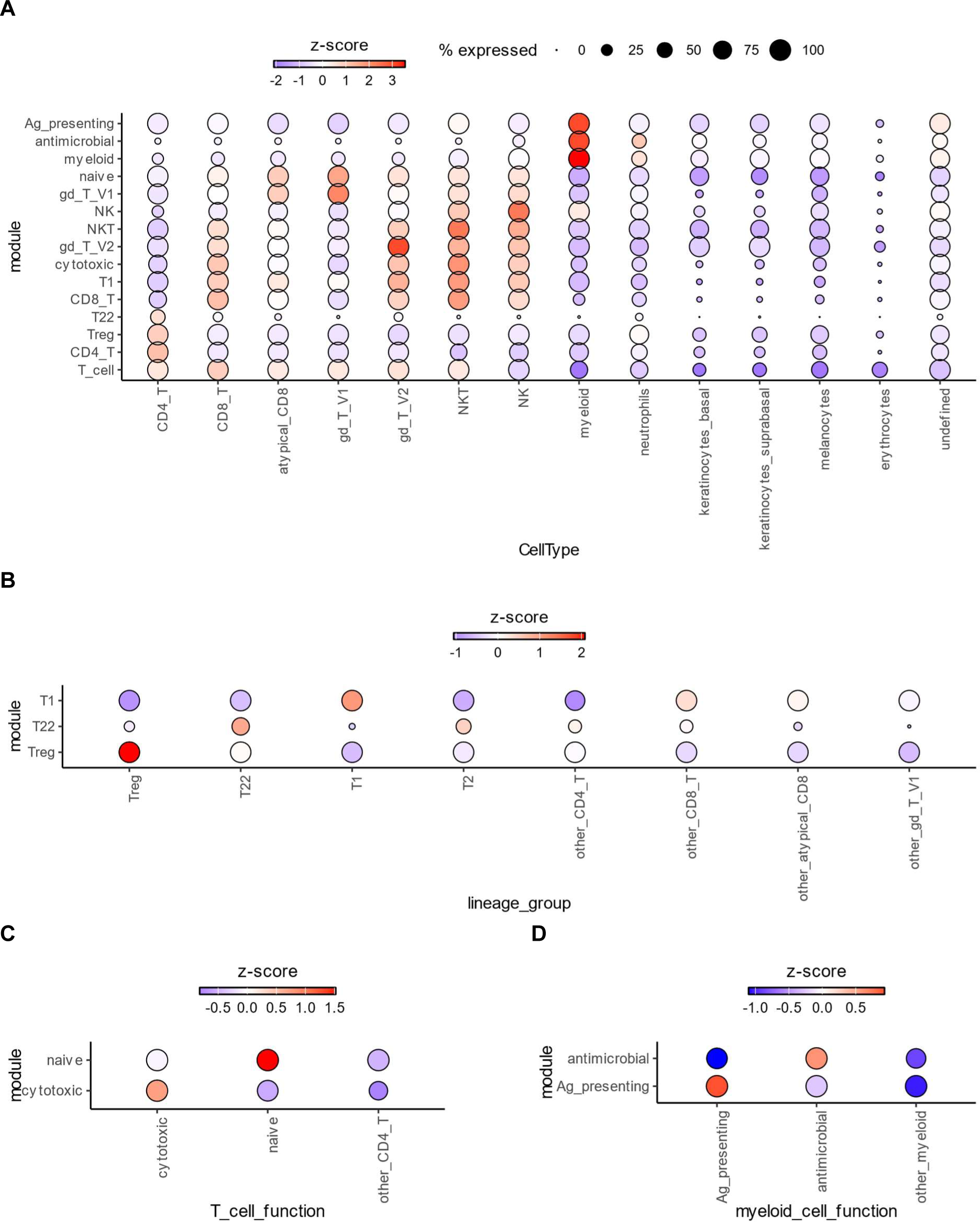
Internal validation of TST blister signatures. Signature Z-scores were calculated for each cell in the single cell RNAseq dataset they were derived from and averaged across all cells annotated with the same cell type or function, as indicated on the x-axis of each dot plot. In the dot plots, signatures are shown on the y-axis, and cell groups are shown on the x-axis. Dot size shows the percentage of cells in each group that expresses any of the signature genes. The size legend is the same for panels A-D. The colour bar indicates the Z-scores and differs for each dot plot. A. Z-scores were calculated compared to all cells in the dataset, to assess differential expression of all derived signatures across all ontogeny cell types. B and C. Z-scores were calculated compared to all T cells in the dataset, to assess differential expression of T cell functional signatures across all T cells. D. Z-scores were calculated compared to all myeloid cells in the dataset, to assess differential expression of myeloid functional signatures across all myeloid cells.

**Table 3.** Evaluation of signature performance. Data are area under the receiver operating characteristic curve (AUROC) values in percentage, with 95% confidence intervals in brackets. AUROC values for module scores were calculated for each signature after assigning cells to a binary class (target cell type or not). Internal validation dataset = day 2 TST blisters; external validation dataset = bronchoalveolar cells from individuals with post-COVID lung disease (24). N/A = target cell type not represented in the dataset.

| Module | Internal validation (full dataset) | Internal validation (T cells only) | Internal validation (myeloid cells only) | External validation |
| --- | --- | --- | --- | --- |
| T_cell | 97.7 (97.6 - 97.8) | N/A | N/A | 99.5 (99.4 - 99.5) |
| NK | 99.6 (99.6 - 99.7) | N/A | N/A | N/A |
| myeloid | 99.9 (99.9 - 99.9) | N/A | N/A | 97.2 (96.9 - 97.4) |
| CD4_T | 95.6 (95.4 - 95.7) | N/A | N/A | 91.1 (90.4 - 91.7) |
| CD8_T | 89.7 (89.5 - 90) | N/A | N/A | 98.2 (98 - 98.4) |
| NKT | 97.5 (97.4 - 97.7) | N/A | N/A | 97.8 (97.4 - 98.2) |
| gd_T_V1 | 95.3 (95 - 95.6) | N/A | N/A | N/A |
| gd_T_V2 | 98.4 (98.1 - 98.7) | N/A | N/A | N/A |
| naive | 93.6 (93.3 - 93.9) | 93.2 (92.9 - 93.6) | N/A | N/A |
| cytotoxic | 84.5 (84.2 - 84.8) | 93.8 (93.6 - 94) | N/A | N/A |
| T1 | 90.3 (90 - 90.5) | 94.2 (94 - 94.4) | N/A | N/A |
| T22 | 78.1 (77.5 - 78.7) | 75.1 (74.5 - 75.8) | N/A | N/A |
| Treg | 99.3 (99.2 - 99.4) | 99.2 (99.1 - 99.3) | N/A | N/A |
| antimicrobial | 99.8 (99.8 - 99.9) | N/A | 97.5 (97 - 97.9) | N/A |
| Ag_presenting | 99.4 (99.3 - 99.5) | N/A | 88.5 (87.1 - 89.9) | N/A |

To validate the signatures in an independent dataset, we made use of our recent single cell analysis of bronchoalveolar cells from individuals with post-COVID lung disease (24). Here, clusters were identified as macrophages, proliferating cells, dendritic cells, CD4 T cells, CD8 T cells, NK T cells, B cells, and two types of lung epithelial cells (club cells and ciliated cells) (24). The TST signatures performed well in this external validation dataset, achieving highest average expression in their target cell type where available, with AUROC values >90 % (**Figure 5**, **Table 3**).

**Figure 5.**
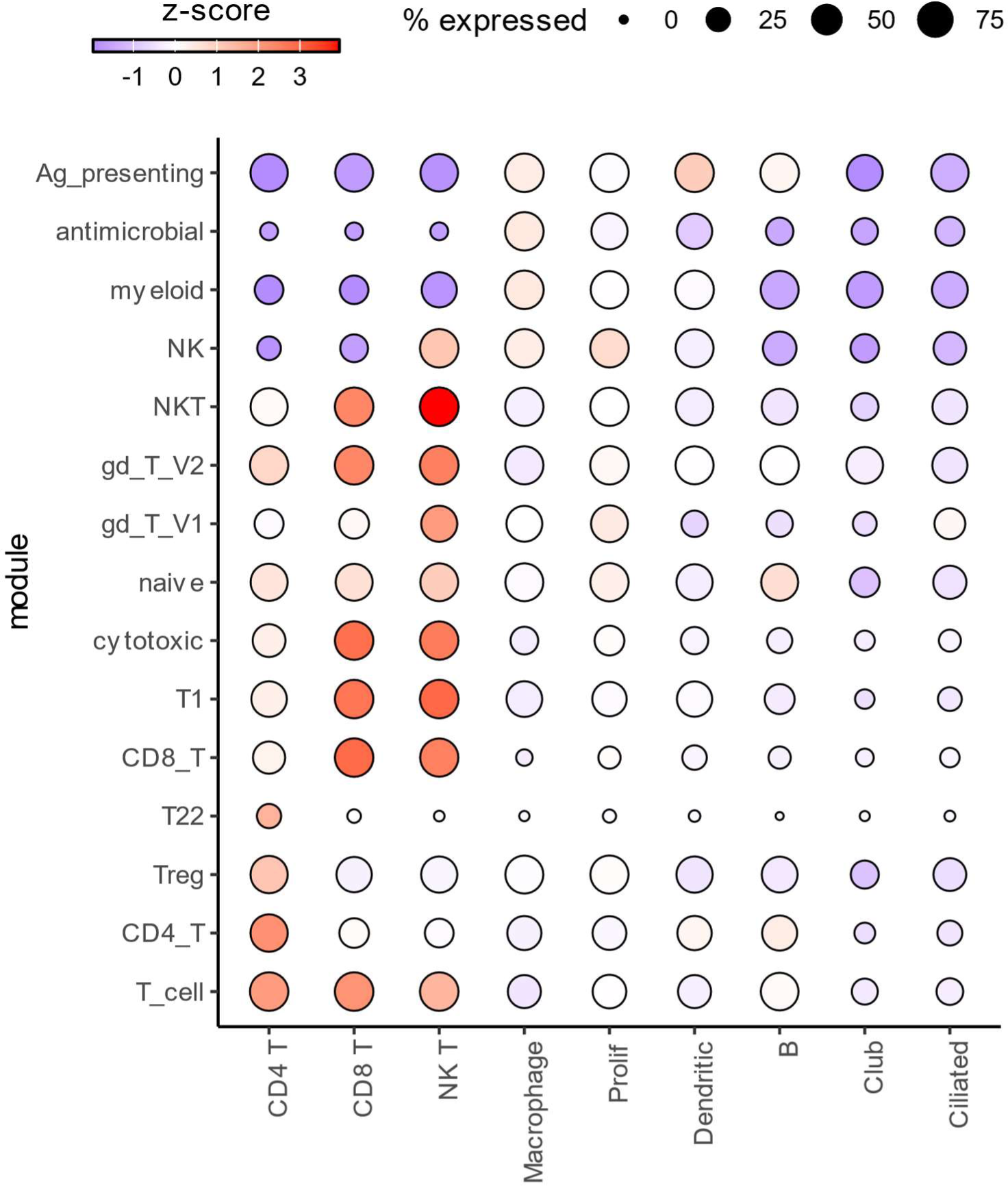
External validation of TST blister signatures in a single-cell RNAseq dataset of bronchoalveolar cells from individuals with post-COVID lung disease (24). Signature Z-scores were calculated for each cell and averaged across all cells annotated with the same cell type, as identified in the original publication. The dot plot shows signatures on the y-axis, and cell type groups on the x-axis. Dot size reflects the percentage of cells in each cell type group that expresses any of the signature genes, while colour indicates the Z-score scaled average expression. Prolif = proliferating cells enriched for macrophage markers.

Taken together, we successfully compiled transcriptional signatures for T cells, myeloid cells and NK cells that are relevant to the immune cell composition of the TST and can discriminate functional T and myeloid cell subsets.

## Discussion

The present study provides the first single cell sequencing characterisation of the human TST. The data are consistent with the existing model of T cell dependent immunity, but also provide novel insights. It is widely accepted that the TST reaction is a Th1 response, amplified by Mtb-specific memory CD4 T cells producing IFNg. However, it has been shown that IFNg levels in TST suction blisters (25) and biopsies (9) are maximal at day 2, whereas IFNg production by CD4 T cells in the TST increases only from day 7 (16,18). We find that the T1 response, detectable on day 2 and manifested by gene expression of IFNG and TBX21, is driven by CD8, gd and NK T cells. Whether these IFNg expressing cells are Mtb-reactive and whether they represent tissue-resident T cells remains of particular interest. We speculate that early IFNg-producing T cells contribute significantly to the subsequent infiltration of memory CD4 T cells. Despite an activated memory phenotype, CD4 T cells in the day 2 TST blisters could not be easily categorised into distinctly polarised T cell lineage sub-populations, such as Th1, Th2 and Th17. A previous study demonstrated that non-specific TCR stimulation of CD4 T cells in vitro leads to an activated phenotype that can be detected at 16 hr, whereas the effect of additional polarising cytokines is only apparent at a later time point (5 days), implying that T cell activation precedes polarisation (26). It is therefore conceivable that T helper lineages are more distinguishable later in the TST. The same in vitro study also found that T cell phenotypes are more distinct after polarisation of naive compared to memory CD4 T cells, emphasising the fluidity of T cell subtypes (26).

There has also been significant interest in type 1 IFN responses to Mtb (23). Whilst we detected the expression of type I IFN-stimulated genes, we were not able to detect any type I IFN gene expression directly. Potential reasons for this may include, the timing of sampling (27), tightly regulated expression levels below the limit of detection in single cell RNA sequencing (28), or sessile behaviour of myeloid producer cells resulting in retention in skin tissue rather than emigration into blister fluid (29). Of note, we found several regulatory cell types in the day 2 TST, including CD4 Treg cells, pDCs and mregDCs. While the presence of Tregs (18) and pDCs (30) in the TST has been described previously, mregDCs are a recently discovered DC population (31) whose role in the TST has not been considered before.

Interestingly, there was a striking absence of B cells in our single cell dataset, in accordance with an independent flow cytometric analysis of TST skin blisters that found 98 % of lymphocytes to be T cells (17). Whilst B cells are not considered to be key players in the TST reaction, they have occasionally been observed in early immunohistochemical studies of the TST (14), and a B cell gene signature is enriched in day 2 TST skin biopsies compared to saline controls, albeit to a lesser extent than other immune cell gene signatures (3,4). The lack of B cells in blister fluid is therefore likely attributable to a reduced ability of B cells to migrate into the blister fluid. A similar observation has been made in a different context, where B cells were not detected in the blister fluid following intradermal injection of antigen-loaded gold nanoparticles, despite being abundant in punch biopsies (32). We also observed only very low numbers of neutrophils in day 2 TST blisters, consistent with a previous report that neutrophils are present in higher numbers at earlier time points (6 hr) during the TST reaction (14). However, neutrophil data are often lacking in tissue single-cell RNAseq analyses, because of their propensity to degrade during sample preparation, low transcriptional activity and subsequent exclusion as presumed empty droplets in the standard CellRanger pipeline (33). An alternative sample processing and analysis strategy might therefore enable better representation of neutrophils in TST blisters.

Finally, we detected a population of naive-like T cells, mapping to Vδ1 T cells and ‘atypical’ CD8 T cells. Whilst the infiltration of antigen-non-specific memory T cells into an inflammatory site has been recognised (34), the migration of naive T cells into non-lymphoid tissue is thought to be restricted (35). There is evidence from a murine model that naive T cells may access non-lymphoid tissue, including the skin, as part of their normal migratory pathway (36). Interestingly, other single cell studies of skin suction blisters have also reported the presence of naive-like CD8 T cells (29,32), though their relevance remains elusive. A predominantly naive phenotype of circulating Vδ1 T cells has been described before (37), and Vδ1 T cells are the main gd T cell subset found in normal skin (38), suggesting that their presence in TST suction blisters might not reflect recruitment from blood. The atypical CD8 and NKT populations in our study were characterised by a comparably low proportion of cells for which we acquired alpha/beta TCR data. This could potentially be the result of a lower expression of the TCR in these cell populations, leading to higher gene drop-out rates. Alternatively, it may be caused by contamination of these clusters with gamma-delta T cells or NK cells, and therefore represent insufficient sub-clustering. In line with this latter interpretation is the presence of alpha/beta TCR sequences in a proportion of cells annotated as gd T cells.

Our study has some limitations. Skin suction blisters may not capture the full cellular response of the TST as they are restricted to mobile cells that migrate out of the skin into the blister fluid upon application of negative pressure. The gene signatures derived here may therefore not fully discriminate against cell types that are not present in TST blisters, such as B cells and neutrophils. In addition, we provide only limited validation of the derived signatures in the discovery dataset itself, and in an independent dataset of bronchoalveolar cells from individuals with post-COVID lung diseases (24). Reassuringly, the TST signatures discriminate successfully between their target cell type and other cell types, including B cells, in the latter. Further external validation in an independently established day 2 TST dataset will yield additional valuable confirmation of the cross-applicability of these signatures in different tissue or disease contexts. Lastly, our study did not include a control group. We can therefore not discriminate between skin-infiltrating cells that responded to the TST challenge as opposed to the general wound process inflicted by skin suction (39).

## Methods

### Study approval

This study was approved by NHS research ethics and UK Health Regulatory Authority (Ref: 18/LO/0680). All study participants provided written informed consent.

### Study cohort and sample collection

Study participants comprised 31 healthy, HIV seronegative adults. All had immune memory to Mtb-specific antigens identified by positive peripheral blood IFNg release assays using the QuantiFERON Gold Plus Test, but no clinical or radiological evidence of active tuberculosis. Participants received intradermal injection of 0.1 ml 2U tuberculin (Serum Statens Institute) in the volar aspect of the forearm as previously described (2–4). At 48 hours, clinical induration at the injection site was measured, before suction blisters were induced at the site of the TST and blister fluid aspirated 2-4 hours later, as described previously (10,16). Erythrocytes were lysed with RBC lysis buffer (Invitrogen).

### Library preparation and sequencing

Up to 20,000 cells per sample (**Table S1**) were stained with the TotalSeq-C Human Universal Cocktail v1 (BioLegend) as per manufacturer’s instructions, and then loaded on to the Chromium controller (10x Genomics) to generate single-cell gel beads in emulsion (GEMs). Single-cell partitioning, reverse transcription, cDNA amplification and library construction were performed using the Chromium Single-cell 5’ Reagent kit v2 (10x Genomics) according to the manufacturer’s instructions to generate gene expression, T cell receptor (TCR) VDJ, and surface protein (= feature barcode) libraries. Libraries were quality checked and quantified using the High Sensitivity D5000kit and 4200 TapeStation System (Agilent) and Qubit 2 Fluorometer (Invitrogen). Sequencing was performed on Illumina’s NovaSeq6000 system, using paired end 150 bp reads, and targeting 20,000 read pairs per cell for gene expression libraries, and 5,000 read pairs per cell for TCR VDJ and feature barcode libraries respectively. Sequencing, sequencing quality control, and conversion of raw data to FASTQ format were undertaken by Novogene Co., Ltd.

### Data processing and analysis

#### CellRanger

Read alignment, feature counting and cell calling was performed with 10x Genomics CellRanger (v7.1.0) against the human genome assembly GRCh38 (gene expression reference version 2020-A and VDJ-T reference version 7.1), using the ‘multi’ pipeline with the ‘expect-cells’ parameter set to 50% of input cells (**Table S1**).

CellRanger’s filtered feature-barcode output matrices were imported to R (version 4.1.1) and combined in a SingleCellExperiment object with the read10xCounts function from the DropletUtils package. Surface protein expression data were stored as ‘alternative Experiment’ inside the same SingleCellExperiment object, allowing isolation of the two modalities, which were processed and utilized independently of each other. Data were processed using several Bioconductor packages, as described below (40).

#### Quality control filtering

Genes with zero counts across all droplets were discarded, resulting in 29761 expressed genes. Doublets were computationally detected and removed with scDblFinder (41). Low-quality cells for each sample, defined as those with comparably low library size, few expressed genes or high number of mitochondrial reads, were identified and removed with the quickPerCellQC function from the scuttle package, which calculates the thresholds for these quality metrics as three median absolute deviations away from the median (42). Lastly, a gene sparsity filter was applied to remove genes that were detected in <0.1% of all cells, yielding 18291 genes. **Table S1** summarizes the number of cell barcodes retained at each step of quality filtering, as well as the quality control thresholds for each sample.

#### Normalisation and integration

Counts were log2 normalised with the deconvolution method implemented in the computePooledFactors function of the scran package (43), before rescaling the size factors between samples with the multiBatchNorm function from the batchelor package (44). Highly variable genes were selected as all those above the mean-variance trendline, using the functions modelGeneVar and getTopHVGs from the scran package, with sample as blocking factor. To adjust for potential batch effects, data from 31 blister samples were integrated with the mutual nearest neighbours algorithm implemented in the fastMNN function of the batchelor package, using the default number of 50 principal components after dimensionality reduction using the highly variable genes.

#### Clustering

Cell clustering was performed with the Louvain algorithm on the ‘corrected’ fastMNN output matrix, which contains the principal component scores for each cell. Louvain clustering was achieved with scran’s clusterCells wrapper function and the SNNGraphParam function from the bluster package. The number of shared nearest neighbours for graph construction was set to k=50, while a range of resolutions was tested using single-cell significance of hierarchical clustering (scSHC) as a post-hoc test to find the resolution that gave the maximum number of statistically confirmed Louvain clusters (20). For the first round of clustering, resolutions between 0.2 and 1 were tested in 0.2 increments. Clusters with at least 500 cells were then sub-clustered a further two times by repeating the workflow of feature selection, data integration and Louvain clustering with scSHC post-hoc test, each time testing resolutions in the range of 0.2 to 3 in 0.2 increments. For the second and third round of clustering, the optimal resolution not only had to yield the maximum number of statistically confirmed Louvain clusters but also be preceded by statistically confirmed clustering at the previous resolution value, unless it was the first resolution that resulted in sub-clusters. Chosen resolutions for each clustering step are summarised in **Table S2**.

#### Feature barcode data

Feature barcode data captured antibody-derived tags (ADT) and allowed quantification of the surface expression for 130 proteins. The DropletUtils and scuttle R packages were used for quality filtering and normalisation of ADT data. The cleanTagCounts function identified and removed cells with low ADT quality. ADT counts were log2-normalised using first the ambientProfileBimodal function to estimate the baseline abundance, and then the medianSizeFactors and logNormCounts functions to calculate size factors and scale ADT counts accordingly.

#### VDJ data

Single cells were identified as TCR-positive if their index barcode was listed in CellRanger’s ‘filtered_contig_annotations’ output files. TCR clones were defined as cells with identical alpha and beta chain CDR3 sequences, as assembled by CellRanger in the ‘cdr3s_aa’ column of the ‘clonotype’ output files. TCR clones found more than once were defined as expanded.

#### Cell type annotation

We compiled a list of hand-curated gene and protein markers based on the *a priori* expectation of cell types and cell states present at the site of the tuberculin skin test (**Table S3**). For each cluster, the percentage of cells in which each marker was detected, as well as the Z-score-scaled average expression of each marker was calculated and visualised in a dot plot. Z-scores were calculated for each cell, either across the entire dataset or across a subset of cells as indicated in the figure legends, and then averaged for each cluster.

To assign ontological and functional annotations to clusters, we iteratively chose selected markers, and visualised their Z-score-scaled expression across a given set of clusters in a heatmap with the pheatmap R package. Hierarchical clustering with ward.D2 linkage was used to gather clusters together based on the similarity of the selected markers, allowing the dendrogram to be split into a desired number of groups. The percentage of TCR-positive cells in a cluster was included as marker, after Z-score-scaling of log2-transformed percentage values.

To identify marker genes for different cell types, the scran R package findMarkers function was implemented using the following settings: test.type = “wilcox”, direction = “up”, pval.type = “all”, lfc = 0. Sample was included as a blocking factor. Genes with an adjusted p-value <0.05 were considered significantly differentially expressed.

#### Gene signatures

Signatures for type 1 and type 2 interferon-stimulated gene expression were described previously (3).

Novel gene signatures were derived from the single-cell data for selected cell types and functions. To make the resulting gene signatures broadly applicable to other datasets, only protein-coding, T cell receptor (TR) and immunoglobulin (IG) genes detected in the scRNAseq data were included. Raw gene counts were summed for each sample-cluster combination, using the aggregateAcrossCells function from the scuttle package, thus creating pseudobulk datasets. The summed counts were normalised by converting them into counts per million and replacing values <0.001 with 0.001 to enable log2 transformation. For each desired signature, the pseudobulk datasets were assigned to a binary class, either the cell type of interest (class 1) or any other cell type (class 0), before 1,000 two-sided Wilcoxon tests were performed for each gene. In each of the 1,000 iterations, a sub-sample of 50 class 1 datasets was compared to an equal-sized sub-sample of class 0 datasets, each randomly selected by sampling with replacement. The final output of the Wilcoxon bootstrapping was the average p-value and average log2 fold difference across all 1,000 iterations. For each signature, significant genes were defined as those up-regulated in the cell type of interest with an adjusted p-value of < 0.05, and up to 50 significant genes were then chosen as signature genes, ranked by decreasing log2 fold difference. To achieve better discrimination, for myeloid function signatures (antimicrobial and Ag-presenting), only genes identified as significant during derivation of the myeloid signatures were selected, and Wilcoxon tests were done exclusively between myeloid clusters, assigned to class 1 or class 0 datasets based on their functional annotation.

To validate the signatures in an external dataset, we utilised our recent analysis of bronchoalveolar cells from individuals with post-COVID lung disease, calculating signature scores across the clusters identified and described before (24).

The signature score for each cell was calculated as the arithmetic mean expression of the constituent signature genes, using log2-normalised counts in the SingleCellExperiment object, and including signature genes with zero expression. For each cluster, the percentage of cells with non-zero signature scores, as well as the Z-score-scaled average signature score, were then calculated and visualised in a dot plot.

To evaluate signature performance, area under the receiver operating characteristic curve (AUROC) values were calculated using the R package pROC. For each signature, cells were assigned to a binary class (target cell type of the module or any other cell type) based on their cluster annotation.

## Data availability

Single cell sequencing data will be available in the ArrayExpress database at the time of peer-reviewed publication of the manuscript.

