## Supplementary Figures for "Single-cell transcriptome and T cell receptor profiling of the tuberculin skin test"

**Figure S1. Repeated sub-clustering separates distinct cell types that are initially grouped together.**

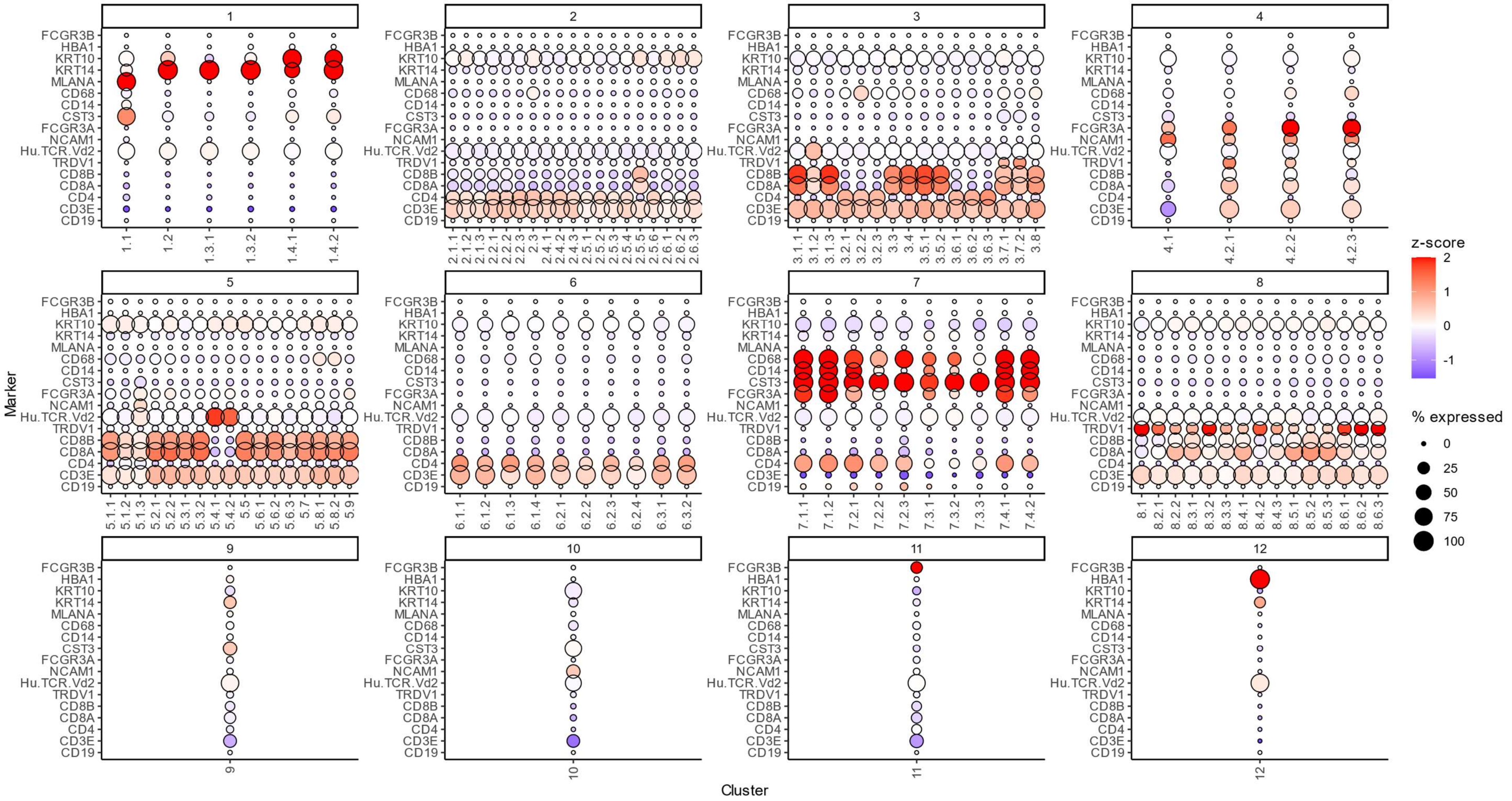


Initial cluster label after a single round of clustering is shown in the box above each dot plot, and final cluster label after two additional rounds of sub-clustering is shown along the x-axis. Dot size represents the percentage of cells expressing each marker in each cluster, and colour shows the Z-score scaled expression of the marker, calculated compared to all other cells in the dataset, and averaged for each cluster. The Z-score colour scale is capped at -2 and 2. Protein markers are prefixed with ‘Hu’.

**Figure S2. Quality metrics of cells retained after quality control filtering, grouped by assigned cell type.**

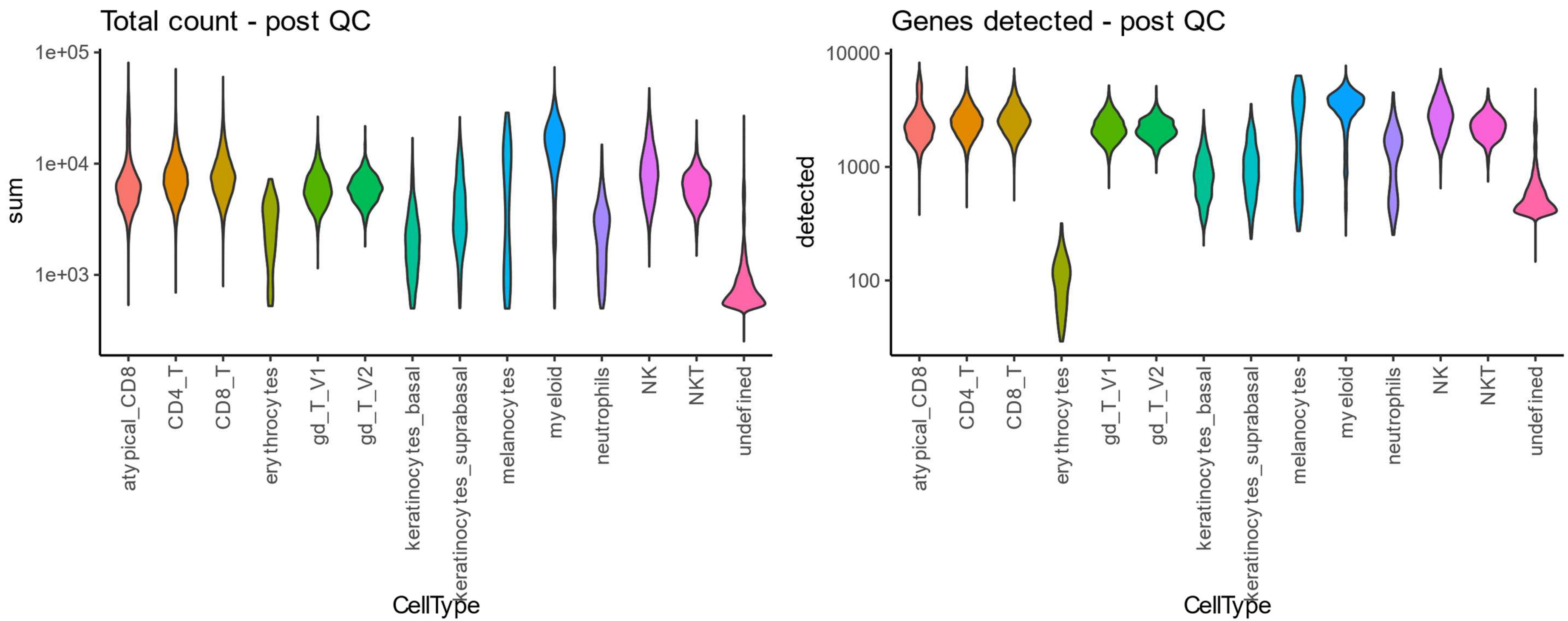


Cells belonging to the ‘undefined’ cluster show a lower number of total unique molecular identifier (UMI) counts (left) and detected genes (right), compared to most other identified cell types.

**Figure S3. Expression of interferon and interferon-stimulated genes.**

**A**

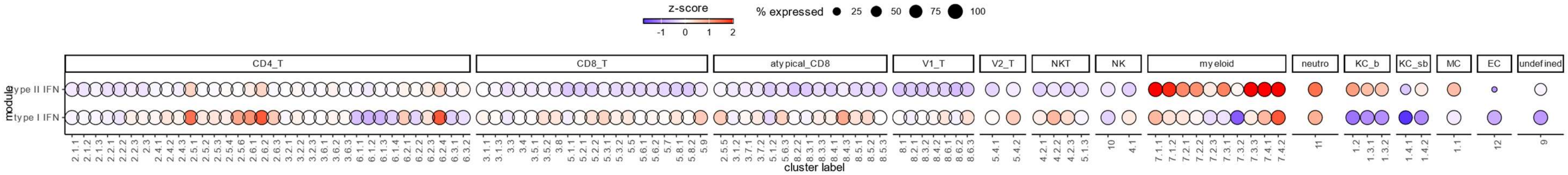


**B**

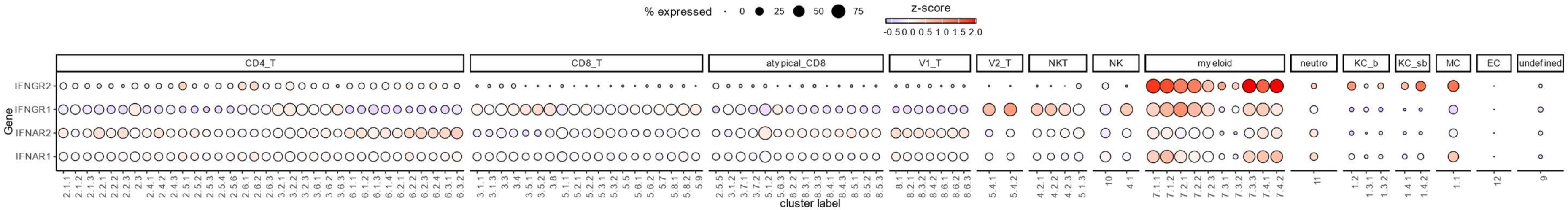


Expression of type I or type II interferon-stimulated multi-gene signatures (**A**) and gene expression of interferon receptors (**B**) in each cluster, stratified by assigned cell type. Dot size represents the percentage of cells expressing each signature or gene in each cluster, and colour shows the Z-score scaled expression of the signature or gene. The Z-score colour scale is capped at -2 and 2. V1_T = Vδ1 gamma/delta T cells; V2_T = Vδ2 gamma/delta T cells; neutro = neutrophil; KC_b = keratinocytes_basal; KC_sb = keratinocytes_suprabasal; MC = melanocytes; EC = erythrocytes

**Figure S4. Internal validation of TST blister signatures.**

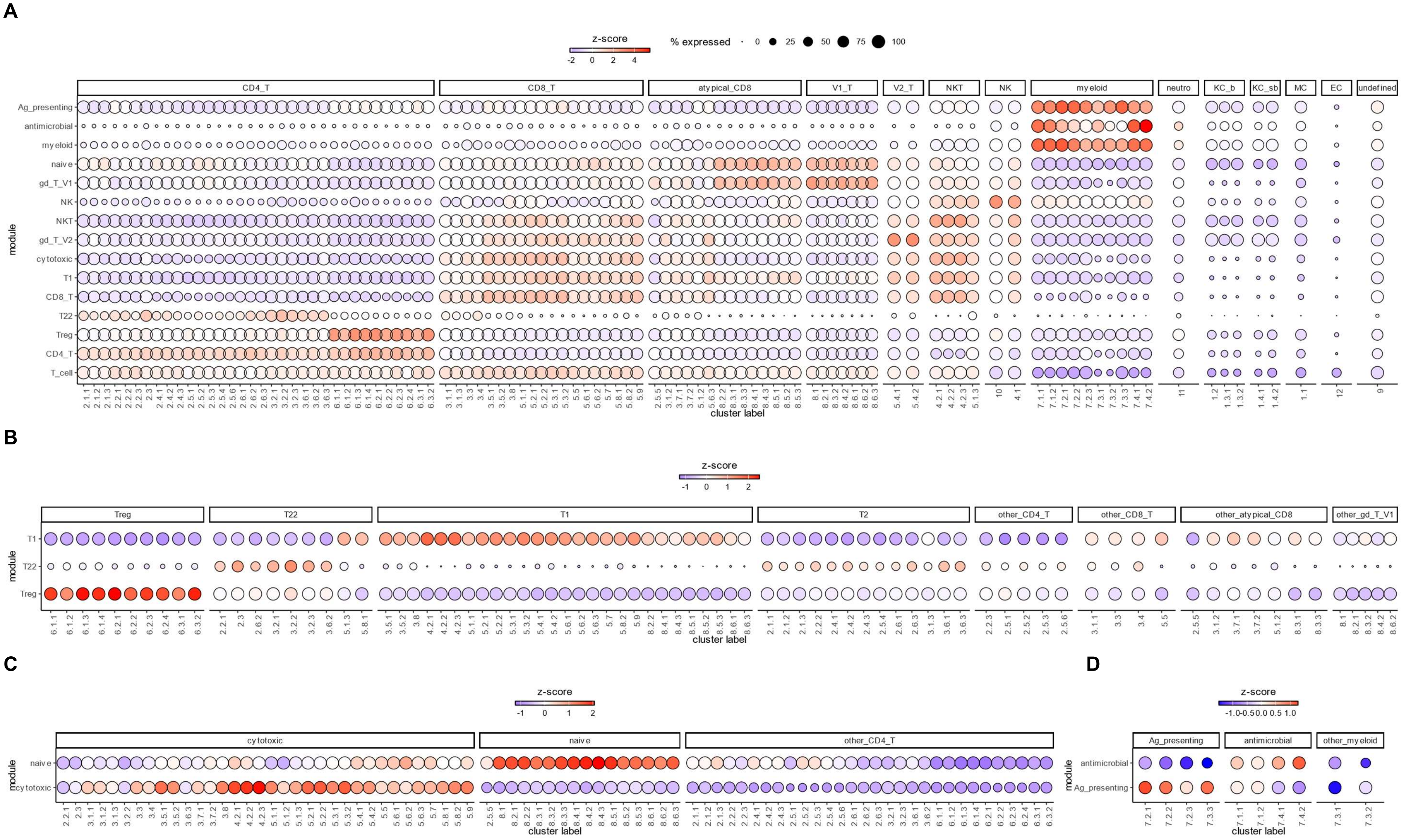

Signature Z-scores were calculated for each cell in the single cell RNAseq dataset they were derived from and averaged across all cells of a given cluster label, as indicated on the x-axis of each dot plot. In the dot plots, signatures are shown on the y-axis, and cluster labels are shown on the x-axis. Dot size shows the percentage of cells in each cluster that expresses any of the signature genes. The size legend is the same for panels A-D. The colour bar indicates the Z-scores and differs for each dot plot. **A.** Z-scores were calculated compared to all cells in the dataset, to assess differential expression of all derived signatures across all ontogeny cell types. **B and C.** Z-scores were calculated compared to all T cells in the dataset, to assess differential expression of T cell functional signatures across all T cells. **D.** Z-scores were calculated compared to all myeloid cells in the dataset, to assess differential expression of myeloid functional signatures across all myeloid cells. V1_T = Vδ1 gamma/delta T cells; V2_T = Vδ2 gamma/delta T cells; neutro = neutrophil; KC_b = keratinocytes_basal; KC_sb = keratinocytes_suprabasal; MC = melanocytes; EC = erythrocytes
